# Effect of progesterone on *Candida albicans* biofilm formation under acidic conditions: a transcriptomic analysis

**DOI:** 10.1101/591347

**Authors:** Bruna Gonçalves, Ruben Bernardo, Can Wang, Nuno A. Pedro, Geraldine Butler, Joana Azeredo, Mariana Henriques, Nuno Pereira Mira, Sónia Silva

**Author notes:** (SS).

## Abstract

Vulvovaginal candidiasis (VVC) caused by *Candida albicans* is a common disease worldwide. A very important *C. albicans* virulence factor is its ability to form biofilms on epithelium and/or on intrauterine devices promoting VVC. It has been shown that VVC has a hormonal dependency and that progesterone affects virulence traits of *C. albicans* cells. To understand how the acidic environment (pH 4) and progesterone (either alone and in combination) modulate *C. albicans* response during formation of biofilm, a transcriptomic analysis was performed together with characterization of the biofilm properties. Compared to planktonic cells, acidic biofilm-cells exhibited major changes in their transcriptome, including modifications in the expression of 286 genes that were not previously associated with biofilm formation in *C. albicans.* The vast majority of the genes up-regulated in the acidic biofilm cells (including those uniquely identified here) are known targets of Sfl1, and the expression of this regulator impaired formation of the acidic biofilm. Under the acidic conditions used, progesterone treatment reduced *C. albicans* biofilm biomass, structural cohesion, matrix quantity and susceptibility to fluconazole. Transcriptomic analysis of progesterone-exposed biofilms led to the identification of 65 down-regulated genes including, among others, the regulator Tec1 and several of its target genes suggesting that the function of this transcription factor is inhibited by the presence of the hormone. Overall, the results of this study show that progesterone modulates *C. albicans* biofilm formation and genomic expression under acidic conditions, which may have implications for *C. albicans* pathogenicity in the vaginal environment.

**Author summary:** Vulvovaginal candidiasis (VVC) is an infection of the vaginal tract that affects millions of women every year. It is caused by fungi of the genus *Candida*, mainly *Candida albicans.* Several *C. albicans* virulence factors contribute to the establishment of this infection, including the ability to form biofilms on vaginal walls and intrauterine devices. *Candida* species belong to vaginal microflora, however under certain conditions they can cause infection. It has been shown that conditions that prompt VVC include those leading to high progesterone levels, as pregnancy. Here we show that progesterone impairs the ability of *C. albicans* cells to form biofilms but causes a potential protective stress response. Indeed, we reveal an increased fluconazole resistance of biofilm cells grown in the presence of the hormone. Additionally, our results suggest that biofilm cells have a specific response to acidic conditions, as those established in the vaginal environment. Deepening the knowledge on the modulation of *C. albicans* virulence by vaginal conditions is essential for a full understanding of the pathogenesis of this species in the vaginal tract and contribute to the disclosure of new targets to treat VVC.

## Introduction

Vulvovaginal candidiasis (VVC) affects millions of women every year and is considered to be an important public health problem. It is estimated that approximately 70-75% of women will experience an episode of VVC in their lifetime [1]. Although VVC is not usually a life-threatening condition, the vaginal tract constitutes a main access route to the bloodstream. Infections of this niche therefore have the potential to result in a severe disseminated infection, particularly in immunocompromised patients [2]. *Candida* species are opportunistic microbes of the commensal microflora, which under certain conditions can transform from symptomless colonization into infection. Most (if not all) women carry *Candida* cells in the vaginal tract at some point in their lives, with or without symptoms of infection [3]. Despite an increased identification of *non-Candida albicans Candida* species (NCAC) [4,5], *Candida albicans* is still the most common species identified in women with VVC [6–8]. The ability of *C. albicans* to form biofilms is an important virulence factor as it confers unique phenotypic characteristics compared to its planktonic counterpart cells, including significant resistance to antifungal agents, host defense mechanisms and physical and chemical stress [9]. In the vaginal environment, *Candida* species can form biofilms on the vaginal epithelium [10] and also on intrauterine devices (IUDs) thereby promoting VVC [11,12]. The ability of *Candida* species to form biofilms in the vaginal environment is an important clinical problem, since the recalcitrant nature of biofilms to currently used antifungals prevents definitive eradication of these microbes from the vaginal lumen thus contributing to the recurrence of VVC [10].

The development of VVC has been associated with the disturbance of the hormonal vaginal environment resulting from behavioral or other host-related factors such as pregnancy, hormone replacement therapy, and use of oral contraceptives or IUDs [1]. It is thought that progesterone contributes to VVC development by stimulating the production of glycogen by epithelial cells [13,14] and inhibiting certain traits of the innate and adaptive immune response [15–17]. Besides these effects on the host it has been also shown that progesterone has direct effects on the physiology of *Candida* cells. Several *Candida* species such as *Candida albicans, Candida guilliermondii, Candida krusei, Candida parapsilosis* and *Candida tropicalis* have corticosterone receptors with high affinity for progesterone [18,19]. A transcriptional survey of the effect of progesterone on *C. albicans* planktonic cells identified activation of stress response pathways, including the induction of genes involved in host immune and drugs response [20]. Banerjee et al. [20] also reported that progesterone decreases *C. albicans* drug susceptibility and found higher MIC for fluconazole and ketoconazole when *C. albicans* planktonic cells are exposed to progesterone. In a previous study, we have shown [21] that progesterone reduces the ability of *C. albicans* strains to form biofilms. This unexpected finding fostered the present work in which we aimed to deepen the current understanding on how progesterone modulates the process of *C. albicans* biofilm formation at the vaginal acidic pH, something that has not been examined before and that is essential for a full understanding of the pathogenesis of this species in the acidic vaginal tract.

## Results and discussion

### Effect of progesterone on *C. albicans* planktonic growth and biofilm formation

To obtain a clearer picture of the effects exerted by progesterone on the physiology of *C. albicans* we examined how the presence of this hormone affects growth of the yeast cells, in either planktonic or biofilm-forming conditions. The assays were performed in RPMI medium at pH 4 in order to mimic the acidic vaginal environment (in the range of 3.6-4.5) [22] and using 2 μM of progesterone, as this is the highest concentration reported in the plasma of pregnant women in the third trimester [21]. In planktonic conditions, the concentration of progesterone used had no significant effect on growth of *C. albicans* SC5314 cells comparatively to the growth curve obtained in unsupplemented RPMI medium (S1 Fig). However, the presence of progesterone reduced the ability of *C. albicans* SC5314 to form biofilms, resulting in a statistically significant decrease on cell cultivability (*p*-value ≤0.0001) and total biomass (*p*-value ≤0.01) (Figs 1A and 1B), compared to biofilms formed in the absence of the hormone. Specifically, we observed a decrease in the number of cultivable cells of approximately 2 orders of magnitude [Log (CFUs/ml)/cm^2^] (Fig 1A) and a decrease in biofilm biomass of approximately 20% (Fig 1B). These results are consistent with those reported by Alves et al. [21]. It is important to stress that the study of Alves et al. [21] was undertaken using two strains (ATCC 90028 and 558234, a reference strain from the American Type Culture Collection (ATCC) and a vaginal isolate, respectively), different from the one used in this study, thereby leading us to conclude that the inhibitory effect of progesterone on biofilm formation by *C. albicans* is independent of the genetic background of the strains used.

**Fig 1.**
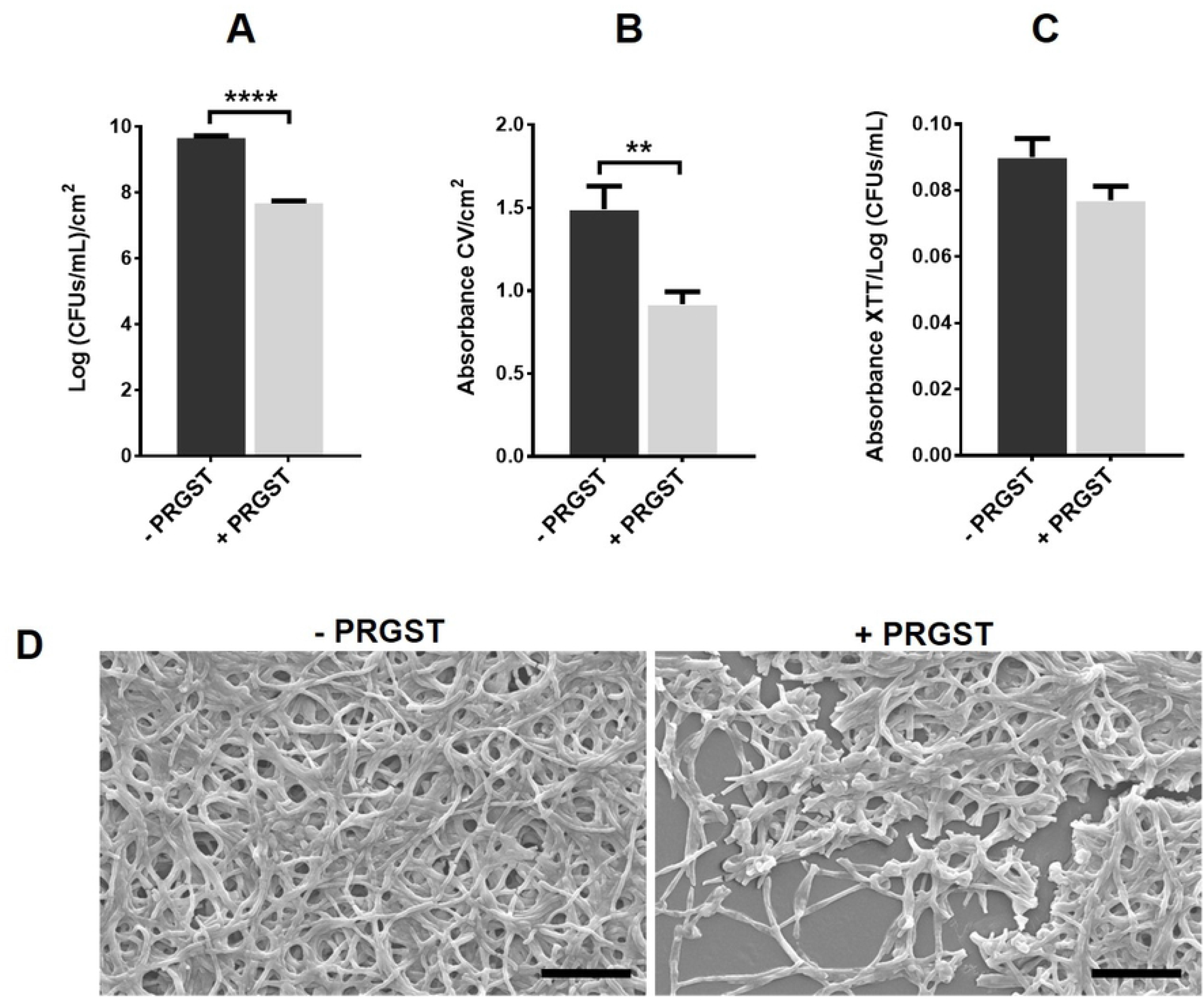
Effect of progesterone on *C. albicans* biofilm formation. **(A)** Cell cultivability determination [Log (CFUs/ml)/cm^2^], **(B)** total biomass quantification (Absorbance CV/cm^2^), **(C)** metabolic activity measurement (Absorbance XTT/Log (CFUs/ml)) and **(D)** scanning electron microscopy images of *C. albicans* SC5314 biofilms grown 24 h in RPMI at pH 4 in absence (-PRGST) or presence of 2 μM of progesterone (+PRGST). Error bars represent standard deviation. Asterisks represent statistical difference between the conditions (**** *p*-value ≥0.0001; ** *p*-value ≥0.01). Original magnification of panel D images was x 1000 and the scale bars correspond to 20 μm.

The effect of progesterone on the metabolic activity of C. *albicans* biofilm’s cells was also evaluated, using XTT reduction assay. The results obtained (Fig 1C) show a slight decrease in the metabolic activity of biofilm cells cultivated in the presence of progesterone, compared to the absence of the hormone; however, this difference is not statistically significant (*p*-value >0.05). The effect exerted by progesterone on the structure and morphological characteristics of the biofilms formed by *C. albicans* was observed by SEM. In the absence of progesterone *C. albicans* SC5314 cells formed a multilayer and compact biofilm that covered the entire surface, constituted by a dense network of hyphal forms (Fig 1D). Differently, in the presence of 2 μM of progesterone, the biofilm consisted of a discontinuous multilayer with a lower number of cells. These observations are consistent with the reduction in the number of cultivable cells (Fig 1A) and total biomass (Fig 1B) of *C. albicans* SC5314 biofilms.

### Effect of progesterone on biofilm matrix production

One of the most important characteristics of *Candida* biofilms is the presence and composition of the extracellular matrix. The functions of the matrix are not entirely clear, but it is thought that it controls the desegregation of the biofilm and protects against antifungal agents and the host immune system [23]. In order to analyze the effect of progesterone on *C. albicans* biofilm matrix production and composition, the total protein and carbohydrate content of the extracellular matrix was determined in the absence and presence of progesterone. Table 1 shows that progesterone led to a significant reduction (*p*-value ≤0.05) of the amount of biofilm matrix to less than half of that formed in its absence. Furthermore, progesterone also led to an alteration of matrix composition, especially the amount of total carbohydrate, which decreases to almost half of the amount detected in the matrix of biofilm formed without progesterone (*p*-value ≤0.05). There is also a slight decrease in the protein level of the biofilm matrix but it was not statistically significant (*p*-value >0.05) (Table 1). Several studies have focused on the identification of matrix components of *C. albicans* biofilms, but very few of them quantified those components [23,24]. Baillie and Douglas [24] reported protein and carbohydrate levels of 5.2% and 41.1% respectively, and Al-Fattani and Douglas [23] reported 5.0% and 39.6%. These values are lower than those obtained in this study for biofilm matrix formed in the absence of progesterone (14.3% and 50.6% for protein and carbohydrate contents, respectively). It should be noted that the growth conditions of the biofilms, including grown medium, pH and *C. albicans* strain, were different in this study and previous reports.

**Table 1.** Progesterone effect on *C. albicans* biofilm matrix composition.

|  | - PRGST | + PRGST |
| --- | --- | --- |
| **Matrix (g/g<sub>biofilm cells</sub>)** | 0.58 ± 0.06 | 0.24 ± 0.02* |
| **Protein (mg/g<sub>matrix</sub>)** | 143.13 ± 10.35 | 120.94 ± 9.27 |
| **Carbohydrate (mg/g<sub>matrix</sub>)** | 505.86 ± 34.03 | 256.84 ± 34.36* |

Matrix quantification (g/g<sub>biofilm cells</sub>) and of its protein (mg/g<sub>matrix</sub>) and carbohydrate (mg/g<sub>matrix</sub>) contents in *C. albicans* SC5314 biofilms grown 24 h in RPMI at pH4, in the absence (- PRGST) or presence of 2μM of progesterone (+PRGST). Asterisks represent statistical difference when compared with biofilm formed in the absence of progesterone (\* *p*-value <0.05).

### Transcriptional profiling of *C. albicans* cells present in biofilms formed in the presence or absence of progesterone

To gain further insights into the molecular mechanisms by which progesterone affects physiology and response of *C. albicans* SC5314 cells, were compared the transcriptomes of biofilms formed in the absence or presence of progesterone (after 24 h of cultivation in RPMI medium at pH4, either supplemented or not with 2 μM of the hormone). Biofilm transcriptomes were also compared with the transcriptome of planktonic *C. albicans* SC5314 cells cultivated for 24h in RPMI growth medium at pH4. With this experimental setup it was possible to characterize the transcriptome of mature biofilms formed by *C. albicans* SC5314 in acidic conditions, something that, to the author’s knowledge, had not been performed before, since previous studies were performed using biofilms formed in near-neutral pH [25–27]. Only genes whose transcripts increased by more than 2-fold in the biofilm cells (in the presence or absence of progesterone), in comparison the expression levels registered in planktonic cells, were considered. A more detailed analysis on the observed transcriptome-wide alterations of *C. albicans* SC5314 biofilm cells follows.

### Transcriptome-wide alterations in *C. albicans* biofilms in the absence of progesterone

Transcriptional profiling of *C. albicans* biofilms formed after 24h of cultivation in acidic RPMI medium (at pH 4) showed a significant (*p*-value below 0.01) alteration (above or below 2-fold) in the expression of 1013 genes, comparing to planktonic cells. Specifically, 616 genes were up-regulated in biofilm cells, while 397 genes were down-regulated. A subset of these genes is listed in Table 2 and the full list is available in S1 Table. About 322 of the up-regulated genes were previously shown to be induced during biofilm formation by *C. albicans* [25–29], however, 294 genes are reported to be up-regulated in biofilm cells for the first time, according to the information available on *Candida* Genome Database (www.candidagenome.org) [30] (highlighted in grey in Table 2 and S1 Table). The identification of these new biofilm-induced genes in our dataset may reflect the acidic pH that was used in this work but not in previous transcriptomic profiling experiments of *C. albicans* biofilms [25–29]. Notably, 8 of the genes found be induced only in our study are essential for normal biofilm formation (highlighted in bold in Table 2 and S1 Table), including *NDT80* and *WOR1*, encoding two transcriptional regulators [31,32]; *ALS5*, encoding an adhesin [33]; the membrane protein *PGA10; CSA2*, encoding a cell surface protein involved in iron utilization [34], and the poorly characterized genes *NPL3, C1_00160C_A* and *C7_00490C_A* [35]. The function of the additional genes with increased expression in the acidic biofilms (286) will require future study.

**Table 2.** Subset of genes found to be biofilm-regulated (up- and down-) in *C. albicans* cells cultivated 24 h at pH 4.

| Up-regulated genes |  |  |
| --- | --- | --- |
| Gene | Biological function | mRNA<sub>biofilm</sub> |
|  |  | mRNA<sub>planktonic</sub> |
| *ALS1* | Cell-surface adhesin | 20.17 |
| *ECE1* | Candidalysin | 18.12 |
| *CAN2* | Basic amino acid permease | 16.91 |
| *SOD5* | Cu-containing superoxide dismutase | 15.01 |
| *C1_13130C_A* | Putative histidine permease | 12.59 |
| *COX3B* | Cytochrome c oxidase involved in mitochondrial respiration | 10.97 |
| *HGT2* | Putative MFS glucose transporter | 10.57 |
| *HYR1* | Hyphal cell wall protein involved in biological adhesion | 9.87 |
| *NAD1* | NADH dehydrogenase involved in mitochondrial respiration | 9.69 |
| *NAD3* | NADH dehydrogenase involved in mitochondrial respiration | 9.49 |
| **C2_10160W_A** | Secreted protein | 9.30 |
| *HGT1* | High-affinity MFS glucose transporter | 9.18 |
| *DIP5* | Dicarboxylic amino acid permease | 8.97 |
| **ATP6** | Subunit 6 of the F0 sector of mitochondrial F1F0 ATP synthase | 7.84 |
| *PRA1* | Cell surface protein that sequesters zinc from host tissue | 5.88 |
| *QDR1* | Putative antibiotic resistance transporter | 5.71 |
| **C7_00490C_A** | Putative AdoMet-dependent proline methyltransferase | 2.93 |
| **NPL3** | Putative RNA-binding protein | 2.63 |
| **CSA2** | Extracellular-associated protein involved in iron assimilation | 2.58 |
| **C1_00160C_A** | Putative nucleolar protein | 2.57 |
| **ALS5** | ALS family adhesin | 2.38 |
| **NDT80** | Meiosis-specific transcription factor | 2.28 |
| **WOR1** | Transcription factor of white-opaque phenotypic switching | 2.16 |

Down-regulated genes
| Gene | Biological function | mRNA<sup>biofilm</sup> |
| --- | --- | --- |
|  |  | mRNA<sup>planktonic</sup> |
| *DUR3* | High affinity spermidine transporter | 0.05 |
| *MEP2* | Ammonium permease | 0.09 |
| *JEN2* | Dicarboxylic acid transporter | 0.11 |
| *C2_00180C_A* | Predicted uricase | 0.15 |
| *C1_01630W_A* | Putative mitochondrial complex I intermediate-associated protein | 0.15 |
| *FDH1* | Formate dehydrogenase | 0.17 |
| *C1_07160C_A* | Protein conserved among the CTG-clade | 0.17 |
| *CWH8* | Putative dolichyl pyrophosphate (Dol-P-P) phosphatase | 0.18 |
| *HSP30* | Putative heat shock protein | 0.18 |
| *LAP3* | Putative aminopeptidase | 0.18 |
| *CTN1* | Carnitine acetyl transferase | 0.18 |
| *C2_01660C_A* | Unknown function | 0.18 |
| *CCP1* | Putative Cytochrome-c peroxidase | 0.18 |
| *MDH1-1* | Predicted malate dehydrogenase precursor | 0.18 |
| *C5_03770C_A* | Putative formate dehydrogenase | 0.18 |
| *C5_03710C_A* | Unknown function | 0.18 |
| *DUR1,2* | Urea amidolyase involved in response to abiotic stimulus | 0.19 |
| *ADH2* | Alcohol dehydrogenase; | 0.19 |
| *PGA13* | GPI-anchored cell wall protein involved in cell wall synthesis | 0.19 |

A subset of genes whose expression was found to increase or decrease (above or below 2-fold) in *C. albicans* SC5314 biofilms grown 24 h in RPMI at pH 4, in comparison with the transcript levels registered in planktonic cells cultivated in the same conditions, was selected and are herein shown, while the full list is available on S1 Table. Genes whose transcription was found to be biofilm-induced or –repressed specifically at the acidic conditions used in this study, are highlighted in grey and among them are highlighted in bold those previously described as essential to biofilm formation. The biological function indicated is based on the information available at Candida Genome Database.

The genes up and down-regulated in the acidic biofilms were functionally clustered using MIPS Functional catalogue (Fig 2). The functional classes enriched (*p*-value below 0.001) in the dataset of up-regulated genes were “transcription”, “protein synthesis”, “protein with binding function” and “transport”. On the other hand, down-regulated genes were enriched in genes involved in “metabolism” (including enrichment of the subclasses “amino acid”, “nitrogen”, “carbohydrate”, “lipid/fatty acid” and “secondary” metabolisms), “generation of energy”, “protein with binding function”, “transport”, “stress response” and “interaction with the environment” (Fig 2). In general, the functional clustering of the genes found to be differently expressed in our acidic biofilms is similar to those found in other studies [25–29] something that could be attributable to the fact that most of the genes that we found differentially expressed in our acidic biofilms have a poorly or even uncharacterized function.

**Fig 2.**
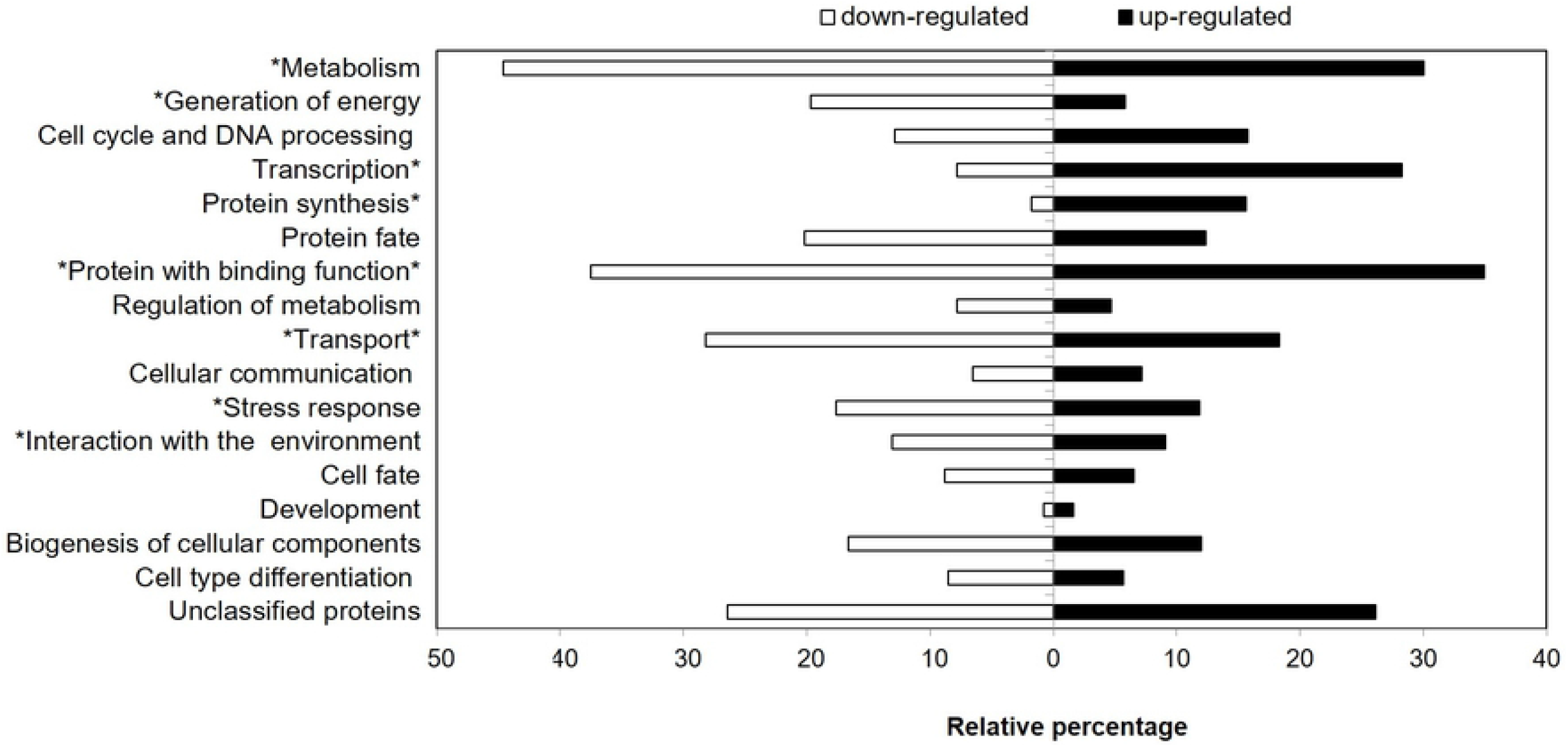
Functional distribution of genes found to be biofilm-regulated in *C. albicans* cells cultivated at pH 4. The genes found to be up- or down-regulated in *C. albicans* SC5314 biofilms grown 24 h in RPMI at pH 4, in comparison with the levels found in planktonic cells cultivated in the same conditions, were clustered according to their biological function, using MIPS Functional Catalogue database (black and white bars correspond to up- and down-regulated genes, respectively). The percentages shown correspond to the number of genes included in each functional class compared to the total number of up- or down-regulated genes. Functional categories considered significantly enriched in the datasets (*p*-value <0.001; taking into consideration the entire genome of *C. albicans* SC5314 reference strain) are indicated with asterisk (asterisk before or after the class denomination means enrichment among down- or up-regulated genes, respectively).

To further understand the transcriptional regulatory network active in the formation of biofilms under acidic conditions we used the PathoYeastract database [36] to cluster the up-regulated genes with transcription factors reported to control the process of biofilm formation in *C. albicans* (Ndt80, Efg1, Bcr1 and Tec1) [31]. Approximately 33% of the genes up-regulated in the acidic biofilms are documented targets of Ndt80 transcription factor, 32% of Efg1, 16% of Bcr1, 13% of Tec1 and 13% of Brg1 (S2 Table). Figure 3 shows the target genes of Efg1, Brg1, Bcr1 and Tec1 with differential expression under biofilm-forming conditions, according with the information available on the PathoYeastract database [36]. A significant overlap between the genes regulated by each transcription factor was observed, confirming the complex and intertwined nature of the regulatory network controlling expression of biofilm genes in *C. albicans* [31]. The high percentage of documented targets of CaNdt80 suggests that this regulator plays a particularly critical role in the control of expression under these conditions. However, 248 upregulated genes are not known targets of Efg1, Ndt80, Bcr1 and Tec1. Analysis using PathoYeastract suggests that 160 of these are targets of Sfl1. In fact, Sfl1 is predicted to regulate 279 genes in the entire dataset, which surpasses the number of targets attributed to Efg1, Ndt80, Bcr1 or Tec1 (Fig 3).

**Fig 3.**
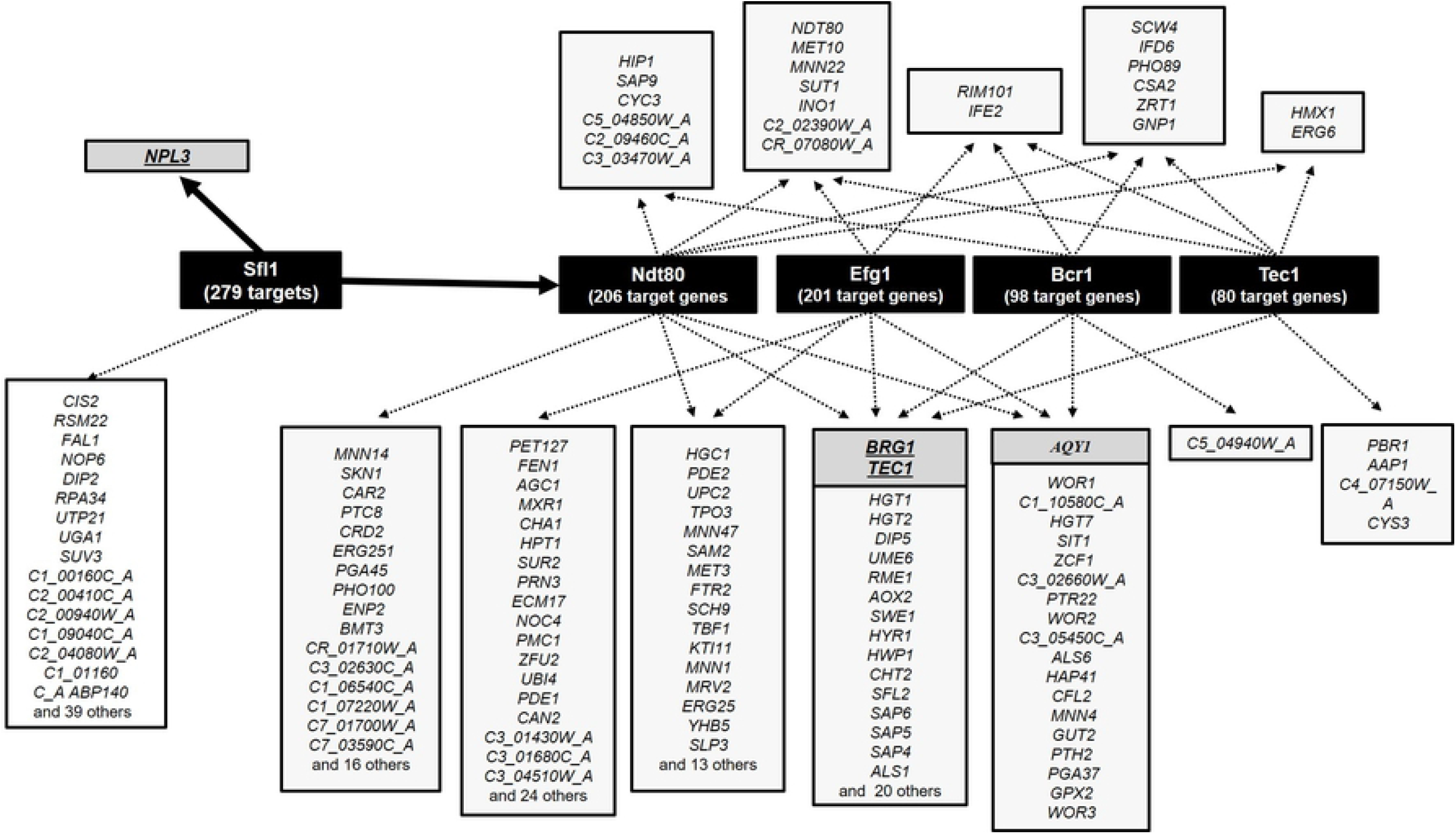
Proposed transcriptional regulatory network underlying the control of *C. albicans* biofilms at pH 4. The genes found to be up-regulated in *C. albicans* SC5314 biofilm cells formed after 24h of cultivation in RPMI at pH 4, in comparison with the levels attained in planktonic cells, were clustered according with the existence of documented regulatory associations with transcription factors mediating the control of transcriptome-wide alterations related with biofilm formation, according with the information available on the PathoYeastract database. The number of documented targets attributed to each transcription factor is indicated in brackets inside the black boxes. Only genes having a biological function related with adhesion and/or biofilm formation (or described to be up-regulated under these conditions) were selected for this analysis, being identified in grey boxes those genes whose deletion was described to abolish biofilm formation. The results that gave rise to this figure are fully detailed in S2 Table.

Deletion of *SFL1* gene almost fully abolished biofilm formation prompted by *C. albicans* cells under acidic conditions (pH 4) (Fig 4A), consistent with this regulator being a critical player in the reprogramming of genomic expression under those conditions. Recently, Sfl1 was found to be required for formation of microcolonies contributing to maximal adhesion to epithelial cells [37]. The involvement of Sfl1 in formation of acidic biofilms represents a further insight into the biological function of this regulator. The high overlap observed between the activated genes in our acidic biofilms documented to be regulated by Sfl1 and Ndt80 prompted us to test whether Sfl1 is a positive regulator of *NDT80* expression. We found that in acidic biofilms formed by the mutant *sfl1Δsfl1Δ* the expression of *NDT80* is approximately 10% of the levels in the parental strain (Fig 4B). Previous studies have also shown that there is functional interaction between Sfl1 and Ndt80 during hyphae formation in *C. albicans* [38]. Sfl1 was also found to be required for maximal expression of *NPL3* (Fig 4B), which is up-regulated in acidic biofilms, and has been shown to be an important determinant of biofilm formation in *C. albicans* [35]. Further studies are required to fully characterize the role of Sfl1 in regulating gene expression during formation of acidic biofilms and, in particular, to identify Sfl1-regulated targets that are essential for biofilm formation. Although Sfl1 has been mostly described as a transcriptional repressor in *C. albicans* [37,39], evidences from chromatin immunoprecipitation profiling have shown that it also acts as a positive regulator [38]. Similarly, the ScSfl1 orthologue has also been found to have a dual effect acting both as a positive and a negative regulator [40]. The molecular mechanisms driving the regulation of the activity of Sfl1 under the different environmental contexts that may control its activity as an activator and/or repressor and how those environmental cues are transduced into that regulatory mechanism remain to be established.

**Fig 4.**
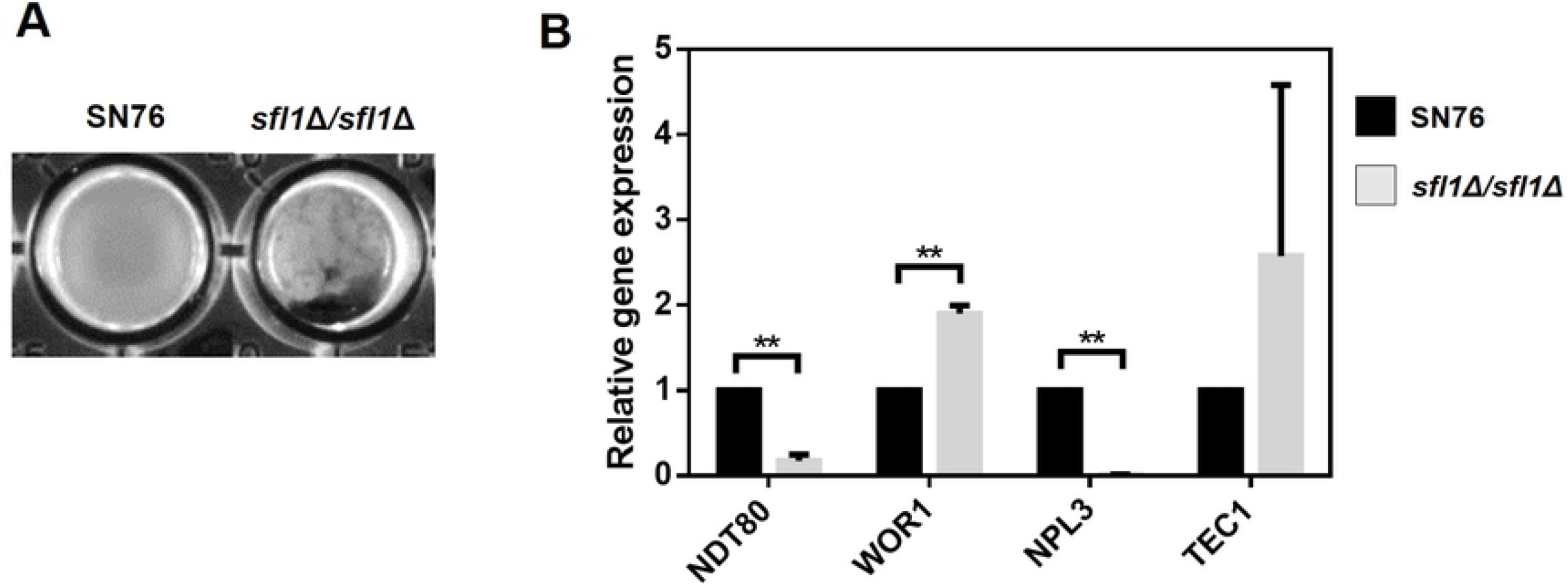
Sfl1 is required for maximal *C. albicans* biofilm formation under acidic conditions (pH 4). **(A)** *Candida albicans* homozygous mutant strain lacking *SFL1 (sfl1Δ/sfl1Δ)* and the respective parental strain (SN76), were cultivated in RPMI at pH 4 for 24h. The biofilms formed under these conditions were photographed and are shown. **(B)** Effect of Sfl1 expression in transcription of *NDT80, WOR1, NPL3* and *TEC1* in biofilm cells after 24h of cultivation in RPMI under the same conditions as those used in the phenotypic assays shown in panel A. Error bars represent standard deviation. Asterisks represent statistical difference between the results of the mutant and parent strain. (** *p*-value ≤0.01).

### Transcriptome-wide alterations in *C. albicans* biofilms in the presence of progesterone

Transcriptome-wide profiling of *C. albicans* biofilms obtained after 24h of cultivation in RPMI medium (at pH 4) supplemented with 2 μM of progesterone identified a significant (*p*-value below 0.01) alteration (above or below 2-fold) in the expression of 1059 genes (553 up- and 503 down-regulated), compared to expression in planktonic cells cultivated in the absence of the hormone (S3 Table). There is a large overlap between this dataset and the dataset of genes differentially expressed in the biofilms formed without progesterone (described above) (S2 Fig). This indicates that most of the changes observed in the progesterone-exposed biofilms result from the process of biofilm formation itself and not from the direct exposure to the hormone. We characterized as progesterone-responsive those genes showing a 40% variation in the biofilms formed in the presence of the hormone, compared to the levels obtained in its absence. Using this criterion, 220 progesterone-responsive genes were identified which were then divided in four groups: *i*) genes up-regulated in biofilms formed in the presence or absence of progesterone, but showing a stronger induction in the presence of the hormone (42 genes); *ii*) genes more strongly up-regulated in biofilms without progesterone than in its presence (87 genes); *iii*) genes more strongly down-regulated in biofilms formed without progesterone (12 genes); *iv*) genes more strongly down-regulated in the progesterone-exposed biofilms (79 genes). A subset of these progesterone-responsive genes is shown in Table 3 and the full list is available in supplementary S4 Table.

**Table 3.** Subset of progesterone-responsive genes in *C. albicans* SC5314 biofilms grown 24 h at pH 4.

| Group *i*) |  |  |  |  |
| --- | --- | --- | --- | --- |
| Gene | Biological function | mRNA<br>+ PRG | mRNA<br>- PRG | $\frac{\text{mRNA} + \text{PRG}}{\text{mRNA} - \text{PRG}}$ |
| *ERG6* | Protein involved in ergosterol biosynthesis | 12.65 | 5.22 | 2.43 |
| *CDR2* | Multidrug transporter of ABC superfamily | 3.15 | 1.35 | 2.33 |
| *PGA31* | Cell wall protein involved in response to drug | 10.18 | 4.85 | 2.10 |
| *PGA45* | Putative GPI-anchored cell wall protein | 20.17 | 9.66 | 2.09 |
| *CDR1* | Multidrug transporter of ABC superfamily | 3.15 | 1.53 | 2.06 |
| *POT1* | Putative peroxisomal 3-oxoacyl CoA thiolase | 2.61 | 1.30 | 2.00 |
| *C3_02870C_A* | Unknown function | 4.74 | 2.49 | 1.90 |
| *GAP2* | General amino acid permease | 2.14 | 1.18 | 1.82 |
| *ATO1* | Putative transmembrane protein | 7.62 | 4.31 | 1.77 |
| *C7_02260W_A* | Unknown function | 3.23 | 1.85 | 1.74 |
| *INO1* | Inositol-1-phosphate synthase | 47.93 | 28.05 | 1.71 |
| *FAS2* | Alpha subunit of fatty-acid synthase | 2.43 | 1.44 | 1.68 |

| *C2_05980C_A* | Putative acyl-CoA hydrolase activity | 3.17 | 1.89 | 1.68 |
| --- | --- | --- | --- | --- |
| *PTH2* | Putative cAMP-independent regulatory protein | 3.60 | 2.19 | 1.64 |
| *AGP3* | Putative serine transporter | 5.12 | 3.20 | 1.60 |
| *C4_00250W_A* | Unknown function | 3.48 | 2.18 | 1.59 |
| *FAT1* | Predicted enzyme of sphingolipid biosynthesis | 2.12 | 1.33 | 1.59 |
| *GIT2* | Putative glycerophosphoinositol permease | 5.04 | 3.19 | 1.58 |
| *CSA2* | Protein involved in iron assimilation | 4.07 | 2.58 | 1.58 |
| **Group ii)** |  |  |  |  |
| **Gene** | **Biological function** | **mRNA**<br>**+ PRG** | **mRNA**<br>**- PRG** | **mRNA + PRG**<br>**mRNA - PRG** |
| *COX3B* | Cytochrome c oxidase | 1.54 | 10.97 | 0.14 |
| *NAD3* | NADH dehydrogenase | 1.38 | 9.49 | 0.15 |
| *ATP9* | ATP synthase required for ATP synthesis | 1.15 | 6.63 | 0.17 |
| *ATP6* | ATP synthase required for ATP synthesis | 1.52 | 7.84 | 0.19 |
| *NAD6* | NADH dehydrogenase | 1.69 | 8.54 | 0.20 |
| *COX2* | Cytochrome c oxidase | 1.76 | 8.56 | 0.21 |
| *WOR3* | Transcription factor | 0.85 | 2.63 | 0.32 |
| *SUT1* | Zn2Cys6 transcription factor | 0.96 | 2.16 | 0.44 |
| *BRG1* | Transcription factor | 0.89 | 2.01 | 0.44 |
| *RME1* | Zinc finger protein; putative meiosis control | 2.20 | 4.55 | 0.48 |
| *PRN3* | Protein similar to pirin; unknown function | 1.53 | 2.84 | 0.54 |
| *PRN1* | Protein similar to pirin; unknown function | 1.61 | 2.90 | 0.56 |
| *TEC1* | TEA/ATTS transcription factor | 3.20 | 5.64 | 0.57 |
| *PGA11* | Putative GPI-anchored protein | 1.16 | 2.03 | 0.57 |
| *TPK1* | cAMP-dependent protein kinase | 2.32 | 3.86 | 0.60 |
| *GIT1* | Glycero phosphodiester transporter | 1.30 | 2.14 | 0.60 |
| *C1_01130W_A* | Putative ubiquitin ligase complex component | 1.99 | 3.24 | 0.61 |
| *WOR1* | Transcription factor of phenotypic switching | 1.32 | 2.16 | 0.61 |
| *IFE2* | Putative alcohol dehydrogenase | 1.50 | 2.41 | 0.62 |
| *SFL2* | Transcription factor required for filament | 1.79 | 2.84 | 0.63 |
| *HAP2* | Transcription factor of low-iron induction | 1.37 | 2.16 | 0.63 |
| *PBR1* | Protein required for biofilm formation | 2.51 | 3.92 | 0.64 |
| *SWE1* | Putative protein kinase required for virulence | 1.36 | 2.11 | 0.64 |
| *AHR1* | Transcription factor involved in adhesion | 1.76 | 2.62 | 0.67 |
| *HYR1* | Hyphal cell wall protein involved in adhesion | 6.82 | 9.87 | 0.69 |

Was selected and listed here a subset of *C. albicans* SC5314 genes found be up-regulated in biofilms grown 24 h in RPMI at pH 4, in the presence (+ PRG) and/or absence (- PRG) of 2 μM of progesterone and whose expression differed more than 40% between the two biofilm-forming conditions. This table includes a subset of genes found to be up-regulated in both conditions, but showing a stronger induction in the presence of the hormone (group *i*) and a subset of genes strongly up-regulated in biofilms without progesterone than on its presence (group *ii*). The full list of progesterone-responsive genes is available on S4 Table. The biological function indicated is based on the information available at Candida Genome Database.

Closer analysis of the set of genes with higher induction in the progesterone-exposed biofilms (clustered in group *i)* revealed a prominent increase in transcription of *CDR1* and *CDR2*, encoding two multidrug resistance transporters of the ABC Superfamily (Table 3). The up-regulation of *CDR1* in response to progesterone in *C. albicans* biofilm cells was further confirmed by qRT-PCR (S3 Fig). Other studies have also reported up-regulation of *CDR1* transcription upon exposure of planktonic *C. albicans* cells to progesterone and to female serum [41,42]. Consistent with this observation, planktonic progesterone-exposed cells are less sensitive to fluconazole than cells cultivated in the absence of the hormone [20,42]. Similarly, we observed that progesterone-exposed biofilm cells were significantly more tolerant to fluconazole than biofilm cells grown in the absence of the hormone (MIC of 1.5 μg/ml compared with 0.25 μg/ml).

The inhibitory effect of progesterone in biofilm formation may be related to reduced expression of genes required for this process. Expression of 166 progesterone-responsive genes is reduced in biofilm cells grown with progesterone (87 clustered in group *ii* and 79 in group *iv* (S4 Table). These include several key regulators of biofilm formation, including the transcription factors Tec1, Brg1, Ahr1, Wor1, Csr1 and Crz2 [31,32,43–45], the heat shock protein Hsp104 [46] and the uncharacterized protein Pbr1 [47], among other genes previously shown to be required for biofilm formation (all highlighted in grey in S4 Table). The progesterone-decreased expression of *TEC1* and *CRZ2* genes in *C. albicans* biofilm cells was further confirmed by qRT-PCR (S3 Fig).

Several genes involved in the process of biofilm formation that had a reduced expression in the presence of progesterone are documented targets of Tec1 (Fig 5A) and Brg1, according to the PathoYeastract database [36]. We therefore tested whether these two regulators are required for biofilm formation in the presence of the hormone. Deleting the *TEC1* reduced the formation of acidic biofilms, which was further aggravated in the presence of progesterone (Fig 5C; data not shown for *BGR1* deletion). Consistently, Tec1 was required for maximal expression of *IFE2, PBR1* and *C7_01510W_A*, either in the presence or absence of progesterone (Fig 5B). Tec1 is also required for maximal expression of *HYR1* in progesterone-exposed biofilm cells. However, in the absence of the hormone expression of *HYR1* is increased in the *tec1Δ/tec1Δ* mutant strain (Fig 5B). This observation is a clear reflection of the complex nature of the regulatory networks governing biofilm formation in *C. albicans*, which can be greatly shaped by the environmental conditions.

**Fig 5.**
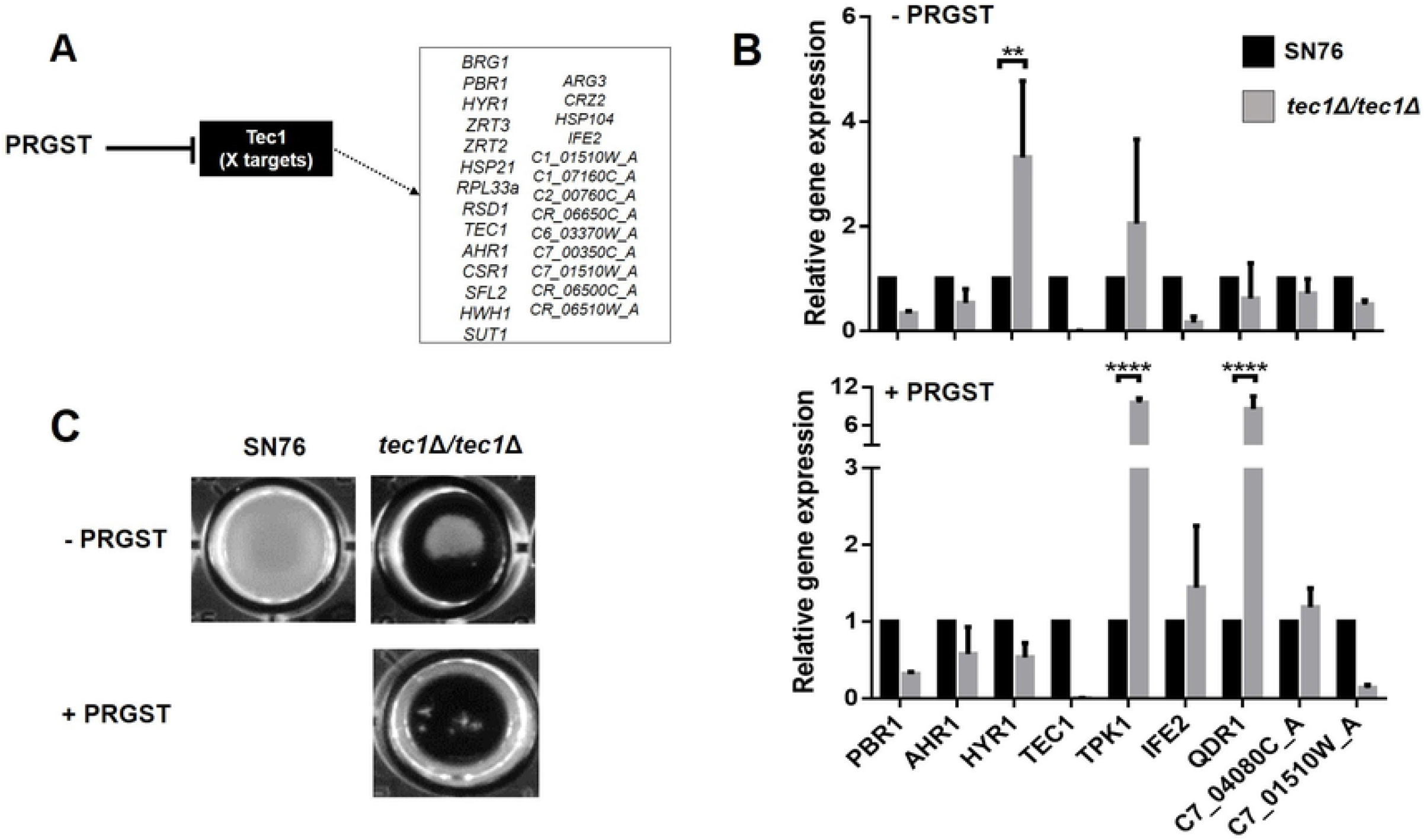
Tec1-documented targets that were found to have a reduced expression in progesterone-exposed biofilm cells. The transcriptome-wide analysis carried out led to the identification of a set of genes (fully detailed in S4 Table) that were found to have a reduced expression in progesterone-exposed biofilm cells, in comparison with the level attained in biofilms not exposed to the hormone. This set of genes included Tec1 and also a set of documented targets. **(A)** In the picture are depicted all those documented Tec1 targets that have a biological function related to biofilm formation or whose expression was found to be induced along biofilm formation. **(B)** The effect of Tec1 in up-regulating genes previously shown as relevant for *C. albicans* biofilm formation was tested by real-time PCR, being confirmed the positive effect in the expression of *PBR1*, *AHR1* and *C7_01510w_A*, both in absence of the hormone (-PRGST) and in progesterone-exposed biofilm cells (+PRGST). Consistent with these results, Tec1 was found to be essential for formation of acidic biofilms and this phenotype was aggravated in the presence of progesterone (C). Error bars represent standard deviation. Asterisks represent statistical difference between the results of the mutant and parent strain.

## Conclusions

Our study examined, for the first time, the alterations occurring in the genomic expression of *C. albicans* during biofilm formation in acidic conditions (RPMI at pH 4), which could potentially lead to the identification of novel players required for maximal biofilm formation in the acidic vaginal tract. Indeed, we identified 286 genes whose transcription was changed during biofilm formation that have not previously been associated with this process. These genes and their regulators represent thus an interesting cohort to search for new players involved in biofilm formation in the acidic vaginal tract. Using this rationale, we showed that Sfl1 is essential for maximal biofilm formation and that is also likely to play a significant role in regulation of gene expression in acidic-biofilm cells. In particular, Sfl1 regulates of Ndt80, a major regulator of biofilm development. The impact of progesterone exposure in biofilm formation and in genomic expression of biofilm-forming *C. albicans* cells was also examined in our work for the first time. The reduced ability to form biofilms observed upon exposure to progesterone was consistent with the reduced transcription of several key genes required for maximal biofilm formation, including the transcriptional regulator Tec1 and several of its target genes. Progesterone exposure was also found to significantly decrease the susceptibility of biofilm cells to fluconazole, which is attributable to a reduction in expression of the drug-efflux pump *CDR1.* Considering that the vaginal tract is one of the main driveways for the development of *C. albicans* infections, the identification of genes that may determine the ability of this yeast to survive in specific vaginal conditions as the acidic pH or progesterone presence may contribute to the disclosure of new targets to treat VVC infections.

## Material and Methods

### Strains and growth conditions

The reference strain *C. albicans* SC5314 was the main strain used in this work. Additionally, some experiments were carried out using *C. albicans* mutant strains and their respective parent. All the strains are listed in Table 4.

**Table 4.** *Candida albicans* strains used in this study.

| Strain name | Parent | Relevant genotype | Reference |
| --- | --- | --- | --- |
| SC5314 |  | Prototrophic | Lab collection |
| SN76 | SC5314 | *arg4Δ/arg4Δ, his1Δ/his1Δ, ura3Δ-iro1Δ::λimm<sup>434</sup>/ura3Δ-iro1Δ::λimm<sup>434</sup>* | [48] |
| *sfl1Δ/sfl1Δ* | SN76 | *sfl1Δ::ARG4/sfl1Δ::HIS1* | [38] |
| *tec1Δ/tec1Δ* | SN76 | *tec1Δ::ARG4/tec1Δ::HIS1* | [38] |
| *brg1Δ/brg1Δ* | SN76 | *brg1Δ::ARG4/brg1Δ::HIS1* | [38] |

For each experiment the strains were subcultured on Sabouraud dextrose agar medium (SDA; Merck, Darmstadt, Germany) for 48 h at 37°C. An inoculum from the SDA plate was suspended in 25 ml of Sabouraud dextrose broth (SDB; Merck, Darmstadt, Germany) and incubated for 18 h under agitation (120 rev/min) at 37°C. Under these conditions the cells were found to be at the beginning of stationary phase. After incubation, the cells were harvested by centrifugation at 3000 g for 10 min at 4°C and washed twice with 15 ml of Phosphate Buffered Saline (PBS; pH 7). Pellets were suspended in Roswell Park Memorial Institute medium (RPMI) (Sigma-Aldrich, St Louis, MO, USA), buffered with MOPS ((3-(N-morpholino) propanesulfonic acid) and adjusted to pH 4 with HCl. The cell density was further adjusted to 1×10^7^ cells/ml for each experiment, using a Neubauer haemocytometer (Marienfeld, Lauda-Königshofen, Germany).

### Effect of progesterone on *C. albicans* planktonic growth

A stock solution of progesterone (Sigma-Aldrich, St Louis, MO, USA) at 10 mM was prepared on ultrapure water and stored at −20°C to be used in all experiments. *C. albicans* SC5314 cells from the pre-inoculum were cultivated in RPMI at pH 4, either supplemented or not with 2 μM of progesterone [21]. Cultures of *C. albicans* cells (1×10^7^ cells/ml with and without progesterone) were placed in 25 ml Erlenmeyer flasks, maintained at 37°C with agitation (120 rev/min) and the increase in optical density at 690 nm was measured over time using a microtiter plate reader (Bio-Tek Synergy HT, Izasa, Winooski, VT, USA). The optical density of the initial cultures was 0.01. The results were presented as optical density over 30h of growth.

### Effect of progesterone on *C. albicans* biofilm formation

#### Biofilm formation

To study the effect of progesterone on biofilm formation, biofilms were developed in presence and absence of progesterone as described by Alves et al. [21]. Briefly, suspensions (1×10^7^ cells/ml) of *C. albicans* SC5314 cells from the pre-inoculum were prepared in RPMI at pH 4, either supplemented or not with 2 μM of progesterone, and placed into wells of 96-wells polystyrene microtiter plates (Orange Scientific, Braine-l‘Alleud, Belgium) (200 μL per well). The plates were incubated for 24 h at 37°C under agitation (120 rev/min). After incubation, the medium was aspirated and non-adherent cells were removed by washing the biofilms with 200 μL of PBS. The analyses of the biofilms were performed in triplicate (biofilms formed from the same pre-inoculum) and in three independent assays (biofilms formed from pre-inoculums independently prepared).

#### Biofilm cultivable cells determination

The number of cultivable cells on biofilms grown with and without progesterone was determined by measuring colony forming units (CFUs) [49]. Briefly, biofilms were scraped from the microtiter plates wells with 200 μL of PBS and the suspensions were vigorously vortexed for 2 min to disaggregate cells. Serial dilutions in PBS of the resuspended biofilms were plated on SDA and incubated for 24 h at 37°C. After incubation, the number of colonies on the SDA plates was counted and the results were presented as total of colony forming units (CFUs) per unit area of microtiter plate well (Log (CFUs/ml)/cm^2^).

#### Biofilm total biomass quantification

Biofilms’ total biomass, formed with and without progesterone, were quantified using crystal violet (CV) staining methodology [49]. Biofilms were fixed on the microtiter plates wells with 200 μL of methanol, which was removed after 15 min. The microtiter plates were allowed to dry at room temperature and biofilms were stained by the addition of 200 μL of CV (1%, v/v) to each well. After 5 min, CV was removed and biofilms were washed twice with sterile water to remove the excess of stain. Finally, 200 μL of acetic acid (33%, v/v) were added to each well to release and dissolve the stain and the absorbance of the obtained solutions was measured in a microtiter plate reader (Bio-Tek Synergy HT, Izasa, Winooski, VT, USA) at 570 nm. The results were presented as absorbance per unit area of microtiter plate well (Absorbance CV/cm^2^).

#### Biofilm metabolic activity determination

A XTT reduction assay [50] was used to determine the metabolic activity of the biofilms formed in the presence and absence of progesterone. A 200 μL aliquot of a solution containing 100 μg/μL of XTT (2, 3-(2-methoxy-4-nitro-5-sulfophenyl)-5-[(phenylamino) carbonyl]-2H-tetrazolium hydroxide) (Sigma-Aldrich, St Louis, MO, USA) and 10 μg/μL of PMS (phenazine methosulfate) (Sigma-Aldrich, St Louis, MO, USA) was added to wells with developed biofilms. The plates were incubated for 3 h in the dark, at 37°C under agitation (120 rev/min). Colorimetric changes were measured at 490 nm using a microtiter plate reader (Bio-Tek Synergy HT, Izasa, Winooski, VT, USA). The absorbance values were normalized with respect to the CFUs and are presented as Absorbance XTT/Log (CFUs/ml).

#### Biofilm structure analysis

The structure of *C. albicans* biofilms formed with and without progesterone was analyzed by scanning electron microscopy (SEM) [21]. Biofilms were formed as described above, except on 24-well polystyrene microtiter plates (Orange Scientific, Braine-l‘Alleud, Belgium) (1 ml of cell suspension per well). Developed biofilms were dehydrated with ethanol (using 70% ethanol for 10 min, 95% ethanol for 10 min and 100% ethanol for 20 min), air dried for 20 min and placed in a desiccator until analysis. Before observation, the base of the wells was removed and mounted onto aluminum stubs, sputter coated with gold. Biofilms were then imaged with an S-360 scanning electron microscope (Leo, MA, Cambridge, USA).

### Effect of progesterone on *C. albicans* biofilm matrix production and composition

In order to study the effect of progesterone on biofilm matrix, biofilms of *C. albicans* SC5314 were formed in the presence and absence of progesterone as described above, using 24-well polystyrene microtiter plates (Orange Scientific, Braine-l’Alleud, Belgium) (1 ml per well). Developed biofilms were scraped from the wells, resuspended in ultra-pure water and sonicated (Ultrasonic Processor, Cole-Parmer, Vernon Hills, IL, USA) for 30 s at 30 W, in order to separate cells from biofilm matrix [49]. Then, the suspensions were vortexed for 2 min and centrifuged at 5000 g for 5 min at 4°C. The matrix-containing supernatants were filtered through a 0.2 μm nitrocellulose filter. The pellets and a portion of supernatants were dried at 37°C until a constant dry weight was obtained. The results of the matrix quantification were presented as matrix dry weight (g) per biofilm’s cells dry weight (g) (g/g_biofilm,cells_).

#### Protein and carbohydrate quantification

The supernatants obtained in the previous step were used to measure protein and carbohydrate contents in the matrices of the biofilms grown with and without progesterone. The protein content was measured using the BCA Kit (Bicinchoninic Acid, Sigma-Aldrich, St Louis, MO, USA) and bovine serum albumin (BSA) as the standard [49]. The results were normalized with the matrix dry weight previously determined and presented as mg of protein per g of matrix dry weight (mg/g_matrix_). Total carbohydrate content was estimated with the phenol-sulfuric method, according to a previous described procedure [51], using glucose as standard. The results were normalized with the matrix dry weight and presented as mg of carbohydrate per g of matrix dry weight (mg/g_matrix_).

### Biofilm cells antifungal susceptibility testing

The susceptibility of *C. albicans* biofilm cells to fluconazole was tested using the reference protocol for broth microdilution antifungal susceptibility testing of yeasts, according to the Clinical and Laboratory Standards Institute methods (NCCLS, M27-A2) [52]. Biofilms were developed in RPMI at pH 4, as described above, in presence and absence of progesterone, using 24-well polystyrene microtiter plates (Orange Scientific, Braine-l‘Alleud, Belgium). Then, biofilms were scraped from the wells with PBS and the suspensions were vigorously vortexed for 2 min and centrifuged at 5000 g for 5 min at 4°C. Pellets were resuspended in RPMI at pH 4 and the cell suspensions were used to the antifungal susceptibility tests and minimum inhibitory concentrations determination (MICs), according to the reference method.

### Transcriptomic analysis

The effect of progesterone on the transcriptome of *C. albicans* biofilms was assessed using species-specific DNA microarrays [53]. For this, the transcriptomes of *C. albicans* SC5314 cells present in biofilms grown in RPMI at pH 4 for 24 h in absence and presence of progesterone (2 μM) were compared with the transcriptome of planktonic cells cultivated for the same time in hormone-free RPMI medium (pH 4). The experimental setups used to cultivate the cells in planktonic and biofilm life styles were the same as described above and using 24-well polystyrene microtiter plates (1 ml per well). Developed biofilms obtained after 24 h of cultivation in presence and absence of progesterone were scraped from the plates with PBS and the suspensions were sonicated (Ultrasonic Processor, Cole-Parmer, IL, USA) for 30 s at 30 W, to separate the cells from the biofilm matrix [49]. Then, all cell suspensions (planktonic and biofilm-forming cells) were centrifuged at 3000 g for 10 min at 4°C, the supernatants were rejected and pellets were used for RNA extraction.

#### RNA extraction

Total RNA extraction was performed using the RiboPure – Yeast Kit (Life Technologies, Carlsbad, CA, USA). *C. albicans* pellets were resuspended in lysis buffer, 10% SDS and a mixture of phenol:chloroform:IAA. The mixtures were vortexed vigorously for 15 s, transferred to screw cap tubes containing cold Zirconia beads and homogenized for 10 min with the vortex set at maximum speed. The obtained lysates were centrifuged for 5 min at 16100 g at room temperature, to separate the aqueous phase, containing the RNA, from the organic phase. Binding buffer and 100% ethanol were added to the aqueous phase and the mixture was applied to filter cartridges assembled in collection tubes and, which were centrifuged for 1 min to pass the mixture through the filter. After that, filters were washed with wash solutions (three times), with centrifugations of 1 min to pass each wash solution through the filter. Then, the filters were centrifuged for 1 min to remove the excess of wash solutions. The filters were transferred to fresh collection tubes and RNA was eluted in two times by applying elution solution (preheated to 95°C) to the filter and centrifuging for 1 min. To remove contaminating chromosomal DNA from the isolated RNA, a DNase digestion reaction was assembled at room temperature (RNA sample, DNase Buffer and DNase I) and incubated for 30 min at 37°C. Then, DNase Inactivation Reagent was added to the mixture and allowed to react for 5 min at room temperature. Lastly, the samples were centrifuged for 3 min at 21000 g to pellet the DNase inactivation reagent and the RNA (supernatant) was transferred to a fresh tube. RNA concentration and purity in each sample was determined by spectrophotometry and integrity was confirmed using an Agilent 2100 Bioanalyzer with an RNA 6000 Nano Assay (Agilent Technologies, Santa Clara, CA, USA).

#### Microarray analysis

cDNA synthesis, hybridization and scanning were performed using protocols similar to those described previously [54], with the exception of that hybridization was carried out using an Agilent hybridization oven at 65°C for 17 h at 100 rpm. In brief, 6 μg of total RNA was incubated with 1.4 μg of anchored Oligo(dT)_20_ primer (Invitrogen, Carlsbad, CA, USA) in a total volume of 18.5 μl for 10 min at 70°C. First-strand buffer (Invitrogen, Carlsbad, CA, USA), 0.5 mMdATP, dTTP, and dGTP; 50 μM dCTP; 10 mM dithiothreitol; 2 μl Superscript III reverse transcriptase (Invitrogen, Carlsbad, CA, USA); and 2 μl Cy3-dCTP or Cy5-dCTP (Amersham, PA53021 and A55021) were added to a total volume of 40 μl, incubated at 42°C for 2 h followed by 1 h at 42°C with an additional 1 μl of Superscript III. RNA was degraded by addition of 1 μl of RNase A at 50 μg/ml and 1 μl of RNase H at 1 unit/μl and incubated at 37°C for 30 min. The labelled cDNAs were purified using a QIAquick PCR purification kit (Qiagen, Hilden, Germany), using a modified protocol. Samples were prepared for hybridization using Agilent’s Two-Color Microarray-Based Gene expression (Quick Amp labelling protocol) and the Gene Expression Hybridization Kit. 10 μL each Cy3-labeled and Cy5-labeled cDNA were used per array (8 × 15 k, design ID 065138), in a total volume of 50 μL. Hybridized microarrays were scanned with an Axon 4000B scanner (Axon Instruments, Union City, CA, USA) and data were acquired with GenePix Pro 5.1 software (Axon Instruments, Union City, CA, USA). Data was analyzed using the LIMMA package in Bioconductor (www.bioconductor.org). Lowess normalization and background correction were applied to each array separately, and quantile normalization was used to allow log ratios to be compared across arrays. Only genes exhibiting log_2_FC >1.0 and having an associated *p*-value below 0.01 were selected for further analysis.

### Measurement of gene transcription based on quantitative real-time PCR (qRT-PCR)

In order to validate some of the results obtained in the microarray analysis, the expression profile of a set of specific genes *(TEC1, CRZ2* and *CDR1)* was obtained using qRT-PCR. The experimental conditions used to cultivate *C. albicans* SC5314 cells and to obtain RNA were the same as those described above for the transcriptomic analysis. Then, 0. 5 μg of total RNA collected from each sample was used to obtain the complementary DNA (cDNA), using the iScript cDNA Synthesis Kit (Bio-Rad, Hercules, CA, USA) according to the manufacturer’s instructions. cDNA synthesis was performed at 70°C for 5 min followed by 42°C for 1 h and lastly 5 min at 95°C to stop the reaction. Approximately 125 ng of the synthesized cDNA were used for the qRT-PCR. qRT-PCR (CF X96 Real-Time PCR System; Bio-Rad, Hercules, CA, USA) was used to determine the relative levels of mRNA transcripts, with the transcript level of *ACT1* mRNA used as an internal control. Primers for the target genes and *ACT1* were designed using Primer3 web software and their sequences are provided S5 Table. Full-length gene sequences were obtained from the *Candida* Genome Database (www.candidagenome.org) [30]. The sequence of each primer was compared to the *Candida* genome database using BLAST [55], to confirm their specificity. The specificity of each primer pair for its corresponding target gene was also confirmed, applying a PCR to genomic DNA extracted from *C. albicans* SC5314 planktonic cells, using the various primer pairs. Then, qRT-PCR was performed using reaction mixtures consisted of SsoFast EvaGreen Supermix (Bio-Rad, Hercules, CA, USA), dH_2_O (Cleaver Scientific Ltd, Warwickshire, UK), cDNA samples and the primer pair respective to the target gene (50 μM). Negative controls (dH2O) and non-reverse transcriptase controls (NRT) were also included in each run. qRT-PCR was performed at 98°C for 2 min in the initial denaturation step, followed by denaturation at 98°C for 5 s and primer annealing at 57°C for 5 s, during 40 cycles. In each cycle it was generated a melting curve running a dissociation stage at 60°C, to verify the amplification product specificity. Control samples were included on each plate to ensure that multiple plates could be compared. The Ct value of each sample was determined and the relative gene expression levels calculated using the ΔCt method [56], being normalized with the internal control gene (Ct_average_= 23.02 ± 1.35). Each reaction was performed in quadruplicate and mean values of relative expression were determined for each gene.

### Experiments using *C. albicans* mutant strains

In order to investigate the role of some genes *(SFL1, TEC1, BRG1)* suggested by our microarray analysis to be important to the biofilm formation in the specific conditions used in this study (pH 4 and presence of progesterone), were formed *C. albicans* biofilms of homozygous null mutants. Mutant and respective parental strains (auxotrophs) used for these experiments are listed in Table 4. Biofilms were developed as previously described for *C. albicans* SC5314 using RPMI at pH 4 supplemented or not with progesterone. Biofilms grown for 24 h were washed with PBS and imaged with ChemiDoc Image System (Bio-Rad, Hercules, CA, USA). Additionally, gene expression of mutants’ biofilm cells was analyzed by qRT-PCR. For that, biofilm formation, RNA extraction and qRT-PCR procedures were performed as previously described for quantitative RT-PCR analyses of *C. albicans* SC5314. Primers for the target genes were designed using Primer3 web software and their sequences are provided in S5 Table.

### Statistical Analysis

Results concerning the determination of biofilm cultivable cells, total biomass, metabolic activity, biofilm matrix quantity and of its components (carbohydrate and protein) and also the results of the qRT-PCR were statistically analyzed using t tests implemented in GraphPad Prism 6 software. All tests were performed with a confidence level of 95 %.

## Acknowledgments

The authors thank Drs. Christophe d’Enfert, Karl Kuchler, Sophie Bachellier-Bassi and Sabrina Jenull for kindly providing mutant strains.

## Supporting information

**S1 Table. List of genes found to be biofilm-regulated (up- and down) in *C. albicans* cells cultivated 24 h at pH 4**. Genes whose expression was found to increase or decrease (above or below 2-fold) in *C. albicans* SC5314biofilms grown 24 h in RPMI at pH 4 in comparison with the transcript levels registered in planktonic cells cultivated in the same conditions, were selected and are herein shown. Genes whose transcription was found to be biofilm-induced or -repressed specifically at the acidic conditions used in this study are highlighted in grey and among them are highlighted in bold those previously described as essential to *C. albicans* biofilm formation. The biological function indicated is based on the information available at Candida Genome Database.

**S2 Table. Regulatory associations between biofilm-induced genes and biofilm-related transcriptions factors**. The genes found to be up-regulated in C. albicans SC5314 biofilms developed 24 h in RPMI at pH 4, in comparison with planktonic cells, were clustered according to the regulatory associations with transcription factors mediating the control of transcriptome-wide alterations related with biofilm formation, based on the information available on the PathoYeastract database. Genes found to be up-regulated specifically at the acidic conditions used in this study are highlighted in grey.

**S3 Table. List of genes found to be altered (up- and down-regulated) in *C. albicans* biofilms grown 24 h in RMPI supplemented with progesterone**. Genes whose expression was found to increase or decrease (above or below 2-fold) in *C. albicans* SC5314 biofilms grown 24 h in RPMI (pH 4) containing 2 μM of progesterone, in comparison with the transcript levels registered in planktonic cells grown in hormone-free medium, were selected and are herein shown. The biological function indicated is based on the information available at Candida Genome Database.

**S4 Table. List of progesterone-responsive genes in *C. albicans* SC5314 biofilms grown 24 h at pH4**. Were selected and listed here *C. albicans* SC5314 genes found to be up- and down-regulated in biofilms grown 24 h in RPMI at pH 4, in the presence (+ PRG) and/or absence (-PRG) of 2 μM of progesterone, and whose expression differed more than 40% between the two biofilm-forming conditions. Progesterone-responsive genes are presented in four groups *i)* genes up-regulated in biofilms formed in the presence or absence of progesterone, but showing a stronger induction in the presence of the hormone (42 genes); *ii)* genes more strongly up-regulated in biofilms without progesterone than on its presence (87 genes); *iii)* genes more strongly down-regulated in biofilms formed without progesterone (12 genes); *iv)* genes more strongly down-regulated in the progesterone-exposed biofilms (79 genes). Progesterone-decreased genes (groups *ii* and *iv)* required for biofilm formation (bold) and/or analyzed by qRT-PCR (Fig 5B) are highlighted in grey. The biological function indicated is based on the information available at Candida Genome Database.

**S5 Table. Forward (FW) and reverse (RV) primers used for qRT-PCR**

**S1 Fig. Progesterone effect on *C. albicans* planktonic cells**. Planktonic growth curves of *C. albicans* SC5314 cells cultivated in RPMI at pH 4 in absence (-PRGST) or presence of 2 μM of progesterone (+PRGST).

**S2 Fig. Venn diagram showing the number of *C. albicans* genes whose transcription was found to be altered (up- or down-regulated) in biofilms formed with and/or without progesterone**. Genes whose expression was found to increase or decrease (above or below 2-fold) in *C. albicans* SC5314 biofilms grown 24 h in RPMI at pH 4, with and/or without 2 μM of progesterone, in comparison with the transcript levels registered in planktonic cells grown in hormone-free medium were selected for this analysis.

**S3 Fig. Transcript levels, estimated by qRT-PCR, of *C. albicans TEC1, CDR1* and *CZR2* genes**. Are presented the levels produced in *C. albicans* SC5314 biofilms cells cultivated 24 h in RPMI at pH 4 in the absence (-PRGST) or presence of 2 μM of progesterone (+PRGST). The values of the transcript levels were normalized using as internal control the levels of ACT1 mRNA. Error bars represent standard deviation. Asterisks represent statistical difference between the conditions (* p-value <0.05; \*\*\**p*-value <0.0001).

## References

1. Sobel JD. Vulvovaginal candidosis. Lancet. 2007 Jun 9;369(9577):1961–71.

2. Ascioglu S, Rex JH, de Pauw B, Bennett JE, Bille J, Crokaert F, et al. Defining opportunistic invasive fungal infections in immunocompromised patients with cancer and hematopoietic stem cell transplants: an international consensus. Clin Infect Dis. 2002 Jan;34(1):7–14.

3. Beigi RH, Meyn LA, Moore DM, Krohn MA, Hillier SL. Vaginal yeast colonization in nonpregnant women: a longitudinal study. Obstet Gynecol. 2004;104(5 Pt 1):926–30.

4. Ahmad A, Khan AU. Prevalence of Candida species and potential risk factors for vulvovaginal candidiasis in Aligarh, India. Eur J Obstet Gynecol Reprod Biol. 2009 May;144(1):68–71.

5. Cetin M, Ocak S, Gungoren A, Ulvi Hakverdi A. Distribution of *Candida* species in women with vulvovaginal symptoms and their association with different ages and contraceptive methods. Scand J Infect Dis. 2007 Jan;39(6–7):584–8.

6. Amouri I, Sellami H, Borji N, Abbes S, Sellami A, Cheikhrouhou F, et al. Epidemiological survey of vulvovaginal candidosis in Sfax, Tunisia. Mycoses. 2011 Sep;54(5):e499–505.

7. Fan SR, Liu XP, Li JW. Clinical characteristics of vulvovaginal candidiasis and antifungal susceptibilities of Candida species isolates among patients in southern China from 2003 to 2006. J Obstet Gynaecol Res. 2008 Aug;34(4):561–6.

8. Vermitsky J-P, Self MJ, Chadwick SG, Trama JP, Adelson ME, Mordechai E, et al. Survey of vaginal-flora *Candidas* pecies isolates from women of different age groups by use of species-specific PCR detection. J Clin Microbiol. 2008 Feb 27;46(4):1501–3.

9. Donlan RM, Costerton JW. Biofilms: survival mechanisms of clinically relevant microorganisms. Clin Microbiol Rev. 2002 Apr 1;15(2):167–93.

10. Harriott MM, Lilly EA, Rodriguez TE, Fidel PL, Noverr MC. Candida albicans forms biofilms on the vaginal mucosa. Microbiology. 2010 Dec;156(Pt 12):3635–44.

11. Chassot F, Negri MFN, Svidzinski AE, Donatti L, Peralta RM, Svidzinski TIE, et al. Can intrauterine contraceptive devices be a *Candida* albicans reservoir? Contraception. 2008 May;77(5):355–339.

12. Lal P, Agarwal V, Pruthi P, Pereira BMJ, Kural MR, Pruthi V. Biofilm formation by Candida albicans isolated from intrauterine devices. Indian J Microbiol. 2008 Dec;48(4):438–44.

13. Dennerstein GJ, Ellis DH. Oestrogen, glycogen and vaginal candidiasis. Aust New Zeal J Obstet Gynaecol. 2001 Aug;41(3):326–8.

14. Špaček J, Buchta V, Jílek P, Förstl M. Clinical aspects and luteal phase assessment in patients with recurrent vulvovaginal candidiasis. Eur J Obstet Gynecol Reprod Biol. 2007 Apr;131(2):198–202.

15. Kalo-Klein A, Witkin SS. Candida albicans: Cellular immune system interactions during different stages of the menstrual cycle. Am J Obstet Gynecol. 1989 Nov;161(5):1132–6.

16. Kalo-Klein A, Witkin SS. Regulation of the immune response to *Candida* albicans monocytes and progesterone. Am J Obstet Gynecol. 1991 May;164(5):1351–4.

17. Keller MJ, Guzman E, Hazrati E, Kasowitz A, Cheshenko N, Wallenstein S, et al. PRO 2000 elicits a decline in genital tract immune mediators without compromising intrinsic antimicrobial activity. AIDS. 2007 Feb 19;21(4):467–76.

18. Loose DS, Stevens D a, Schurman DJ, Feldman D. Distribution of a corticosteroid-binding protein in *Candida* and other fungal genera. J Gen Microbiol. 1983;129(8):2379’2385.

19. Skowronski R, Feldman D. Characterization of an estrogen-binding protein in the yeast *Candida albicans*. Endocrinology. 1989 Apr;124(4):1965–72.

20. Banerjee D, Martin N, Nandi S, Shukla S, Dominguez A, Mukhopadhyay G, et al. A genome-wide steroid response study of the major human fungal pathogen *Candida albicans*. Mycopathologia. 2007 Jun 22;164(1):1–17.

21. Alves CT, Silva S, Pereira L, Williams DW, Azeredo J, Henriques M. Effect of progesterone on *Candida albicans* vaginal pathogenicity. Int J Med Microbiol. 2014 Nov;304(8):1011–7.

22. Boskey ER. Origins of vaginal acidity: high D/L lactate ratio is consistent with bacteria being the primary source. Hum Reprod. 2001 Sep 1;16(9):1809–13.

23. Al-Fattani MA, Douglas LJ. Biofilm matrix of *Candida* albicans and *Candida* tropicalis: chemical composition and role in drug resistance. J Med Microbiol. 2006 Aug;55(Pt 8):999–1008.

24. Baillie GS, Douglas LJ. Matrix polymers of *Candida* biofilms and their possible role in biofilm resistance to antifungal agents. J Antimicrob Chemother. 2000 Sep 1;46(3):397–403.

25. García-Sánchez S, Aubert S, Iraqui I, Janbon G, Ghigo J-M, D’Enfert C. Candida albicans biofilms: a developmental state associated with specific and stable gene expression patterns. Eukaryot Cell. 2004 Apr;3(2):536–45.

26. Murillo LA, Newport G, Lan C-Y, Habelitz S, Dungan J, Agabian NM. Genome-wide transcription profiling of the early phase of biofilm formation by *Candida albicans*. Eukaryot Cell. 2005 Sep; 4(9):1562–73.

27. Yeater KM, Chandra J, Cheng G, Mukherjee PK, Zhao X, Rodriguez-Zas SL, et al. Temporal analysis of *Candida albicans* gene expression during biofilm development. Microbiology. 2007 Aug 1;153(8):2373–85.

28. Desai J V, Bruno VM, Ganguly S, Stamper RJ, Mitchell KF, Solis N, et al. Regulatory role of glycerol in *Candida albicans* biofilm formation. MBio. 2013 Jan 9;4(2):e00637–12.

29. Nett JE, Lepak AJ, Marchillo K, Andes DR. Time course global gene expression analysis of an in vivo *Candida biofilm*. J Infect Dis. 2009 Jul 15;200(2):307–13.

30. Skrzypek MS, Arnaud MB, Costanzo MC, Inglis DO, Shah P, Binkley G, et al. New tools at the *Candida* Genome Database: biochemical pathways and full-text literature search. Nucleic Acids Res. 2010 Jan;38(Database issue):D428–432.

31. Nobile CJ, Fox EP, Nett JE, Sorrells TR, Mitrovich QM, Hernday AD, et al. A recently evolved transcriptional network controls biofilm development in *Candida albicans*. Cell. 2012 Jan 20;148(1-2):126–38.

32. Yi S, Sahni N, Daniels KJ, Lu KL, Huang G, Srikantha T, et al. Self-induction of a/a or alpha/alpha biofilms in *Candida albicans* is a pheromone-based paracrine system requiring switching. Eukaryot Cell. 2011 Jun;10(6):753–60.

33. Garcia MC, Lee JT, Ramsook CB, Alsteens D, Dufrêne YF, Lipke PN. A role for amyloid in cell aggregation and biofilm formation. PLoS One. 2011 Mar 8;6(3):e17632.

34. Srikantha T, Daniels KJ, Pujol C, Kim E, Soll DR. Identification of genes upregulated by the transcription factor Bcr1 that are involved in impermeability, impenetrability, and drug resistance of *Candida albicans* a/α biofilms. Eukaryot Cell. 2013 Jun;12(6):875–88.

35. Bonhomme J, Chauvel M, Goyard S, Roux P, Rossignol T, d’Enfert C. Contribution of the glycolytic flux and hypoxia adaptation to efficient biofilm formation by *Candida* albicans. Mol Microbiol. 2011 May;80(4):995–1013.

36. Monteiro PT, Pais P, Costa C, Manna S, Sá-Correia I, Teixeira MC. The PathoYeastract database: an information system for the analysis of gene and genomic transcription regulation in pathogenic yeasts. Nucleic Acids Res. 2017 Jan 4;45(D1):D597–603.

37. McCall AD, Kumar R, Edgerton M. Candida albicans Sfl1/Sfl2 regulatory network drives the formation of pathogenic microcolonies. PLoS Pathog. 2018 Sep 25;14(9):e1007316.

38. Znaidi S, Nesseir A, Chauvel M, Rossignol T, D’Enfert C. A comprehensive functional portrait of two heat shock factor-type transcriptional regulators involved in *Candida albicans* morphogenesis and virulence. PLoS Pathog. 2013 Aug 15;9(8):e1003519.

39. Bauer J, Wendland J. Candida albicans Sfl1 suppresses flocculation and filamentation. Eukaryot Cell. 2007 Oct;6(10):1736–44.

40. Conlan RS, Tzamarias D. Sfl1 functions via the co-repressor Ssn6-Tup1 and the cAMP-dependent protein kinase Tpk2. J Mol Biol. 2001 Jun 22;309(5):1007–15.

41. Karnani N, Gaur NA, Jha S, Puri N, Krishnamurthy S, Goswami SK, et al. SRE1 and SRE2 are two specific steroid-responsive modules of *Candida* drug resistance gene 1(CDR1) promoter. Yeast. 2004 Feb;21(3):219–39.

42. Larsen B, Anderson S, Brockman A, Essmann M, Schmidt M. Key physiological differences in *Candida albicans CDR1* induction by steroid hormones and antifungal drugs. Yeast. 2006 Aug;23(11):795–802.

43. Finkel JS, Xu W, Huang D, Hill EM, Desai J V, Woolford CA, et al. Portrait of *Candida albicans* adherence regulators. PLoS Pathog. 2012 Feb;8(2):e1002525.

44. Askew C, Sellam A, Epp E, Mallick J, Hogues H, Mullick A, et al. The zinc cluster transcription factor Ahr1p directs Mcm1p regulation of *Candida albicans* adhesion. Mol Microbiol. 2011 Feb;79(4):940–53.

45. Nobile CJ, Nett JE, Hernday AD, Homann OR, Deneault J-S, Nantel A, et al. Biofilm matrix regulation by *Candida albicans* Zap1. PLoS Biol. 2009 Jun;7(6):e1000133.

46. Fiori A, Kucharikova S, Govaert G, Cammue BPA, Thevissen K, Van Dijck P. The heat-induced molecular disaggregase Hsp104 of *Candida albicans* plays a role in biofilm formation and pathogenicity in a worm infection model. Eukaryot Cell. 2012 Aug;11(8):1012–20.

47. Sahni N, Yi S, Daniels KJ, Srikantha T, Pujol C, Soll DR. Genes selectively up-regulated by pheromone in white cells are involved in biofilm formation in *Candida albicans*. PLoS Pathog. 2009 Oct;5(10):e1000601.

48. Noble SM, Johnson AD. Strains and strategies for large-scale gene deletion studies of the diploid human fungal pathogen *Candida albicans*. Eukaryot Cell. 2005 Feb;4(2):298–309.

49. Silva S, Henriques M, Martins A, Oliveira R, Williams D, Azeredo J. Biofilms of non-Candida albicans Candida species: quantification, structure and matrix composition. Med Mycol. 2009 Nov 3;47(7):681–9.

50. Hawser S. Comparisons of the susceptibilities of planktonic and adherent *Candida albicans* to antifungal agents: a modified XTT tetrazolium assay using synchronised *C*. albicans cells. Med Mycol. 1996 Jan 1;34(2):149–52.

51. DuBois M, Gilles KA, Hamilton JK, Rebers PA, Smith F. Colorimetric method for determination of sugars and related substances. Anal Chem. 1956 Mar;28(3):350–6.

52. NCCLS. Reference Method for Broth Dilution Antifungal Susceptibility Testing of Yeasts; Approved Standard — Second Edition. NCCLS Doc M27-A2. 2002;22(15).

53. Bernardo RT, Cunha D V., Wang C, Pereira L, Silva S, Salazar SB, et al. The CgHaa1-regulon mediates response and tolerance to acetic acid stress in the human pathogen *Candida glabrata*. Genes|Genomes|Genetics. 2017 Jan 5;7(1):1–18.

54. Rossignol T, Logue ME, Reynolds K, Grenon M, Lowndes NF, Butler G. Transcriptional response of *Candida parapsilosis* following exposure to farnesol. Antimicrob Agents Chemother. 2007 Apr 30;51(7):2304–12.

55. Altschul S, Madden TL, Schäffer AA, Zhang J, Zhang Z, Miller W, et al. Gapped BLAST and PSI-BLAST: a new generation of protein database search programs. Nucleic Acids Res. 1997 Sep 1;25(17):3389–402.

56. Livak KJ, Schmittgen TD. Analysis of relative gene expression data using real-time quantitative PCR and the 2(-Delta Delta C(T)) Method. Methods. 2001 Dec;25(4):402–8.

